## Supporting Information for "Application of sequential cyclic compression on cancer cells in a flexible microdevice"

### Supplementary Figures

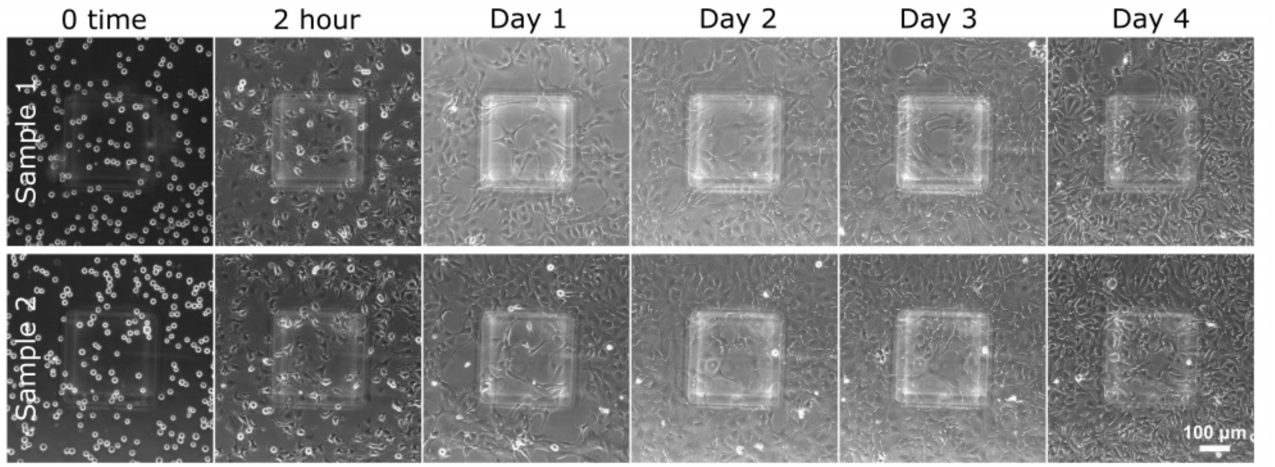


**S1 Fig. On-chip growth of SKOV-3 cells on Matrigel-coated glass surface in micro-piston device.** Representative images of two samples during culture from zero time to Day 4 after piston-retracted loading of cells at -615 mbar.

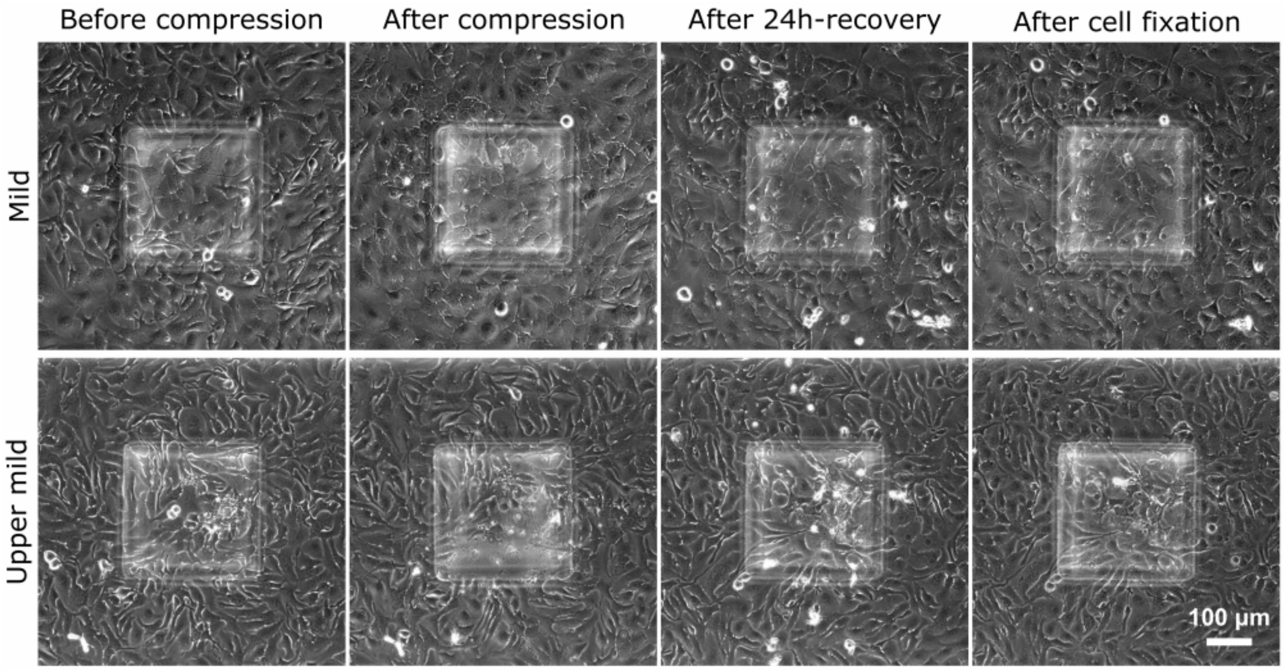


**S2 Fig. Examples of 24h-recovery of mildly and upper-mildly compressed SKOV-3 cells on Matrigel-coated glass surface in micro-piston device.**

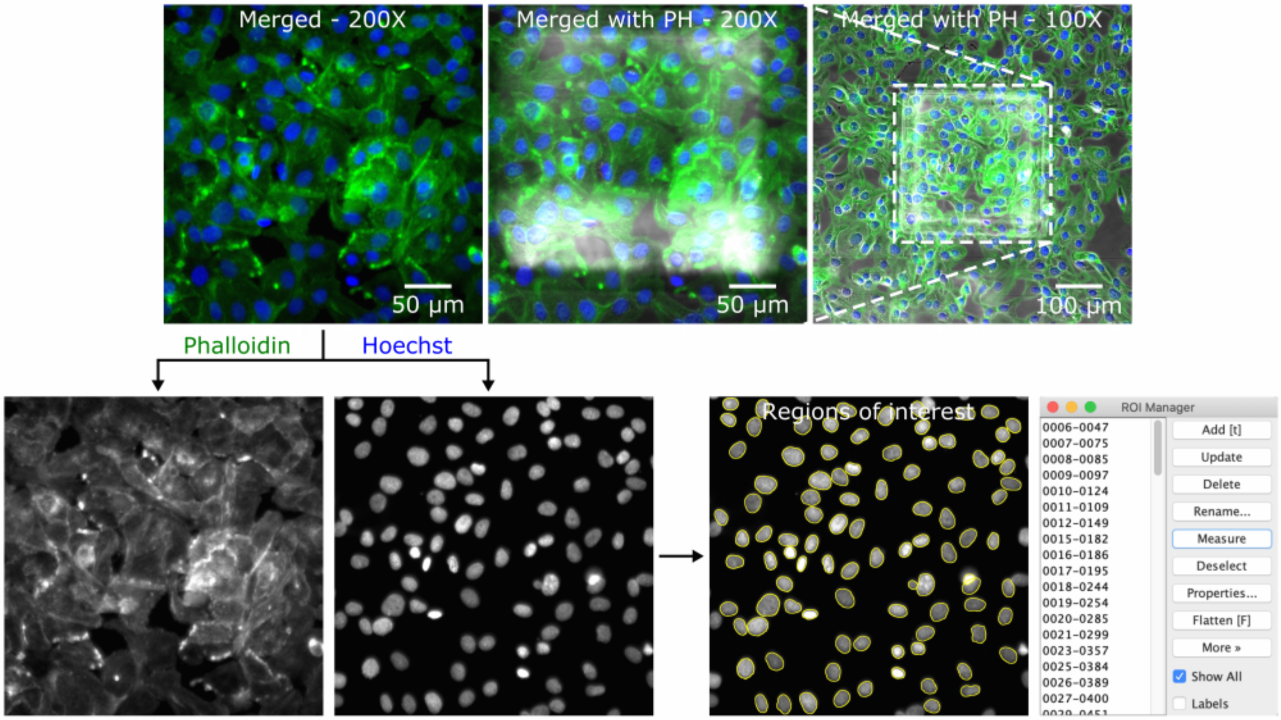


**S3 Fig. Image analysis of nuclear shape after cell compression on Matrigel-coated surfaces**. Regions of interest (ROIs) extracted for the cell nuclear boundaries in Hoechst (stain for nuclei) epi-fluorescence channel (blue in merged/composite frames) were used to measure area, circularity, and aspect ratio of the cell nuclei, as independent of phalloidin (stain for actin) epi-fluorescence channel (green in merged/composite frames). Merged - 200X: merged form of the phalloidin and Hoechst epi-fluorescence images obtained at two-hundred-fold magnification; merged with PH - 200X: merged form of two-hundred-fold magnification epi-fluorescence images with the corresponding phase-contrast (PH) image for the region under micro-piston; merged with PH - 100X: merged form of one-hundred-fold magnification epi-fluorescence images with the corresponding phase-contrast (PH) image for the region under and around micro-piston in channel.

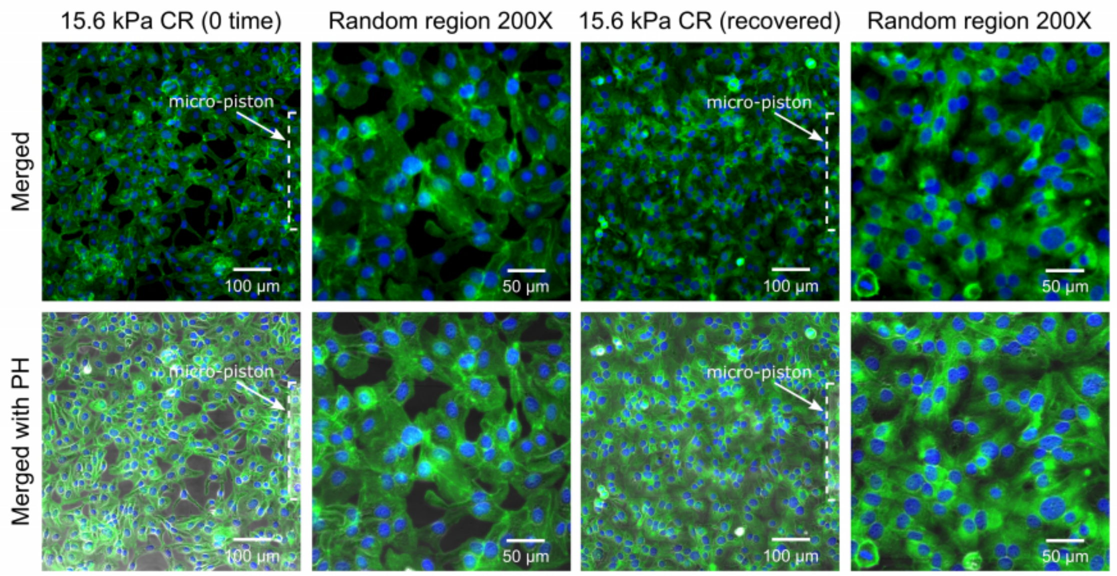


**S4 Fig. End point assay staining and analysis for actin and nuclei of cancer cells in control region (CR) fixed at zero time or after 24 h-recovery, following 1 hour-long cyclic compression between 0 and 15.6 kPa.** Control cell groups stained for actin (green) and nuclei (blue) for their form at zero time and 24 h-recovery. Merged: merged form of the phalloidin (stain for actin) and Hoechst (stain for nuclei) epi-fluorescence images; merged with PH: merged form of the epi-fluorescence images with the corresponding phase-contrast (PH) image; Random region 200X: two-hundred-fold magnification images of control cells in random region as part of the control region.

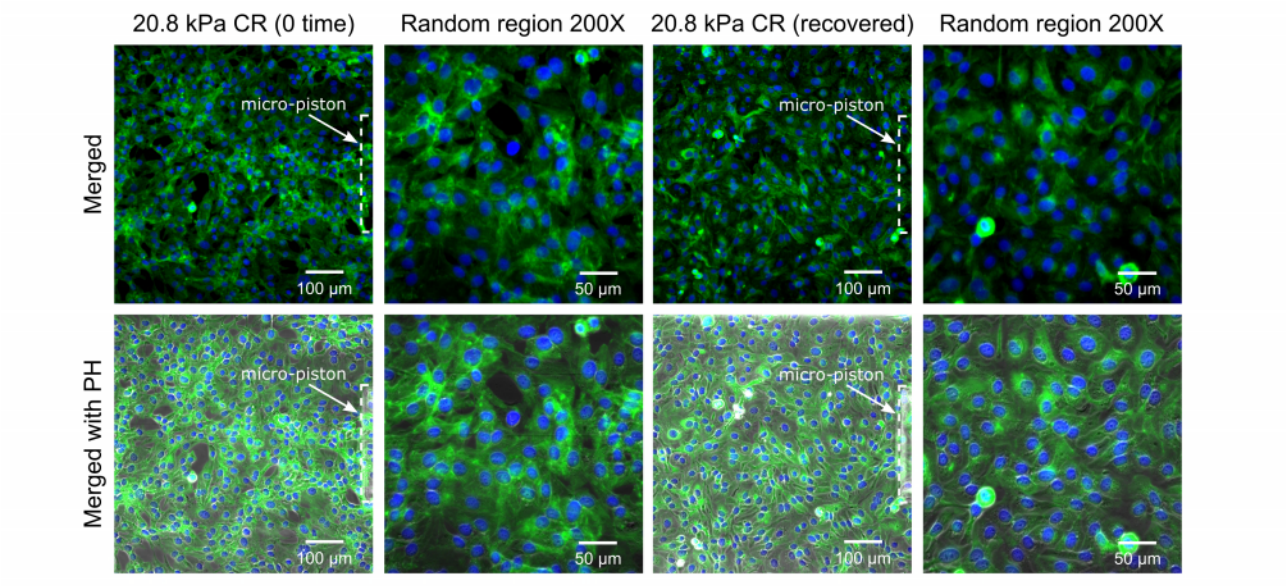


**S5 Fig. End point assay staining and analysis for actin and nuclei of cancer cells in control region (CR) fixed at zero time or after 24 h-recovery, following 1 hour-long cyclic compression between 0 and 20.8 kPa.** Control cell groups stained for actin (green) and nuclei (blue) for their form at zero time and 24 h-recovery. Merged: merged form of the phalloidin (stain for actin) and Hoechst (stain for nuclei) epi-fluorescence images; merged with PH: merged form of the epi-fluorescence images with the corresponding phase-contrast (PH) image; Random region 200X: two-hundred-fold magnification images of control cells in random region as part of the control region.

### Supplementary Movies

**S1 Movie. Temporal evolution of the dynamic cell compression within Matrigel-coated micro-piston devices.** Time-lapse live cell imaging with 100 ms per frame was recorded for ladder pressure increase from 0 kPa up to 15.6 kPa and a short cycle at 15.6 kPa with gradual increase (from 0 kPa to 15.6 kPa) and decrease (from 15.6 kPa to 0 kPa) in 30 s for each. These steps were sequentially followed by 1 h-long cyclic compression between 0 kPa and 15.6 kPa (piston contact pressure from simulation). Cells were compressed in the temporal evolution of the dynamic pressure control within Matrigel-coated micro-piston devices and dynamic changes in cells during the cyclic compression were captured in detail at a rate of 100 ms per frame.

**S2 Movie. Representative data recording of the dynamic changes of GFP-tagged live cell actin during application of sequential cyclic compression at various applied pressures.** SKOV-3 ovarian cancer cells were transduced on-chip with CellLight Actin-GFP, BacMam 2.0 baculovirus (Invitrogen) to study the dynamic biomechanical processes under compression. The actin cytoskeleton profile was quantified for the cancer cell response under applied forces at the compressed stages and at rest stages during the compression process.

### Supplementary Dataset

**S1 File. Corrected total cell fluorescence (CTCF) data for actin integrated density and statistical test results.**
